# Spatial neglect after stroke is reduced when lying inside a 3T MRI scanner

**DOI:** 10.1101/2022.08.01.502290

**Authors:** Hans-Otto Karnath, Hannah Rosenzopf, Britta Stammler, Stefan Smaczny, Axel Lindner

## Abstract

Recently, it was discovered that the static magnetic field of an MRI scanner not only causes horizontal vestibular nystagmus in healthy individuals but, in addition, leads to a horizontal bias of spatial orienting and exploration that closely resembles the one observed in stroke patients with spatial neglect (a disorder of spatial attention and exploration). The present study asked whether the behavioral effects of this magnetic vestibular stimulation (MVS) can be inverted and thus be used to reduce the pathological bias of stroke patients with spatial neglect. Indeed, when patients with left-sided spatial neglect entered the scanner with their feet first, i.e., with the magnetic field vector pointing from head to toes, MVS inside the scanner reduced the ipsilesionally biased distribution of overt attention and the corresponding neglect of the left parts of the search-space. Thus, an intervention as simple as lying in a 3T MRI scanner bears the potential to become an integral part of a future strategy for treating spatial neglect.

## Introduction

While the widespread use of magnetic resonance imaging in the clinical and scientific fields emerged in the 1980s, it has been reported only recently that healthy subjects exposed to the static magnetic fields of MRI scanners exhibit a persistent horizontal vestibular nystagmus. Since this effect’s initial description by Marcelli and co-workers (2009) and subsequent systematic investigation by Roberts and colleagues (2011), several studies have replicated this effect and confirmed the hypothesis that the stimulation of the vestibular organ is induced by Lorentz forces resulting from the interaction of the MR scanner’s static magnetic field with ionic currents in the endolymph fluid of the subject’s labyrinth (see Ward et al., 2019 for review). Only recently, Lindner and co-workers (2021) found that the MR static magnetic field not only causes horizontal vestibular nystagmus in healthy individuals but in addition leads to a horizontal bias of spatial orienting and exploration that closely resembles that seen in stroke patients with spatial neglect.

Spatial neglect is a visuo-spatial attention disorder occurring after stroke, most typically of the right hemisphere. Patients with spatial neglect show spontaneous and sustained deviation of their eyes and head (Fruhmann-Berger & Karnath 2005) as well as focus of attention towards the ipsilesional, right side of space (Heilman et al., 1983; Karnath & Rorden, 2012; Karnath, 2015). When the eye movements of such patients are recorded during visual search, an asymmetric, spatially biased exploratory behavior becomes visible (Karnath et al., 1998), leading to neglect of the contralesional side of the surrounding scene.

The present study was motivated by the aforementioned observation that magnetic vestibular stimulation (MVS) by lying inside a 3T MRI scanner induces spatial biases in healthy subjects that mimic those observed in neglect patients (Lindner et al., 2021). In the main experiment by Lindner and colleagues, the subjects lay in the scanner in the conventional neurological position, namely entering the scanner with their heads first, resulting in the magnetic field vector of our scanner pointing from subjects’ toes to their heads. Under this condition, the center of healthy subjects’ visual scan-path during visual search deviated towards the right, as is also the case for most patients with spatial neglect (compare above). The authors further observed that for an “inverted” positioning in the scanner, i.e., when subjects entered the scanner with their feet first, MVS led to inverted behavioral effects: healthy subjects now showed a bias to the left instead of to the right side. We thus asked whether the behavioral effects of MVS on spatial attention and orientation during “inverse positioning” could be used to reduce the pathological core symptoms of stroke patients with spatial neglect of the left side of space. We examined whether an inverted MVS-effect would directly counteract the pathological bias of patients suffering from spatial neglect.

## Methods

### Participants

Neurological patients consecutively admitted to the Center of Neurology at Tübingen University were screened for manifest spatial neglect. Patients with a left hemispheric stroke, with diffuse or bilateral brain lesions, with lesions restricted to the brainstem or cerebellum, with tumors, with no visible demarcations, or without acute imaging data were not enrolled. A group of three neurological patients suffering from right-sided brain lesions and spatial neglect could be included. Two of these patients exhibited pathological scores in all four neglect tests that were conducted as part of the diagnostic procedure on the day of the experiment (see below); one patient obtained results indicating spatial neglect in three of the four diagnostical tests (Tab. 1).

**Table 1.** Demographic and clinical data of the right brain-damaged patients with spatial neglect.

| **Patient** | **Sex** | **Age** | **Aetiology** | **Time**<br>**since**<br>**lesion** | **CoC**<br>**(letter**<br>**cancel.)** | **CoC**<br>**(bells**<br>**cancel.)** | **EWB**<br>**(line**<br>**bisection)** | **Copying** |
| --- | --- | --- | --- | --- | --- | --- | --- | --- |
| **W.E.** | M | 64 | Isch. | 7 | -0.021 | 0.163* | 0.14* | 2 points* |
| **P.R.** | M | 35 | TBI | 736 | 0.342* | 0.323* | 0.71* | 3 points* |
| **E.B.** | M | 68 | Isch &<br>haem. | 104 | 0.185* | 0.173* | 0.44* | 3 points* |

M, males; Isch., ischemia; haem., haemorrhage; TBI, traumatic brain injury; Time since lesion, days between onset of brain lesion and participation in the experiment; CoC, Center of Cancellation (Rorden & Karnath, 2010); EWB, endpoint weightings bias (McIntosh et al., 2017); \*, values that exceed the cut-offs for the letter and bells cancellation, the line bisection, and the copying tasks, indicating spatial neglect.

Patient W.E. is a 64-year-old right-handed male patient who suffered a stroke 7 days before the present experimental investigation. W.E was admitted to our department after having developed left-sided hemiparesis, dysarthria, and spatial neglect. The initial CT perfusion and CT angiography uncovered a fronto-laterally located perfusion deficit and occlusion of the M2 segment of the middle cerebral artery in the right hemisphere. Lysis therapy was initiated but aborted when the patient indicated it could have been a wake-up stroke (after he had originally documented that symptoms had started while being awake). A magnetic resonance T2 fluid attenuated inversion recovery (FLAIR) sequence one day after admission uncovered a demarcation in right frontal and insular cortices (Fig. 1).

**Fig. 1.**
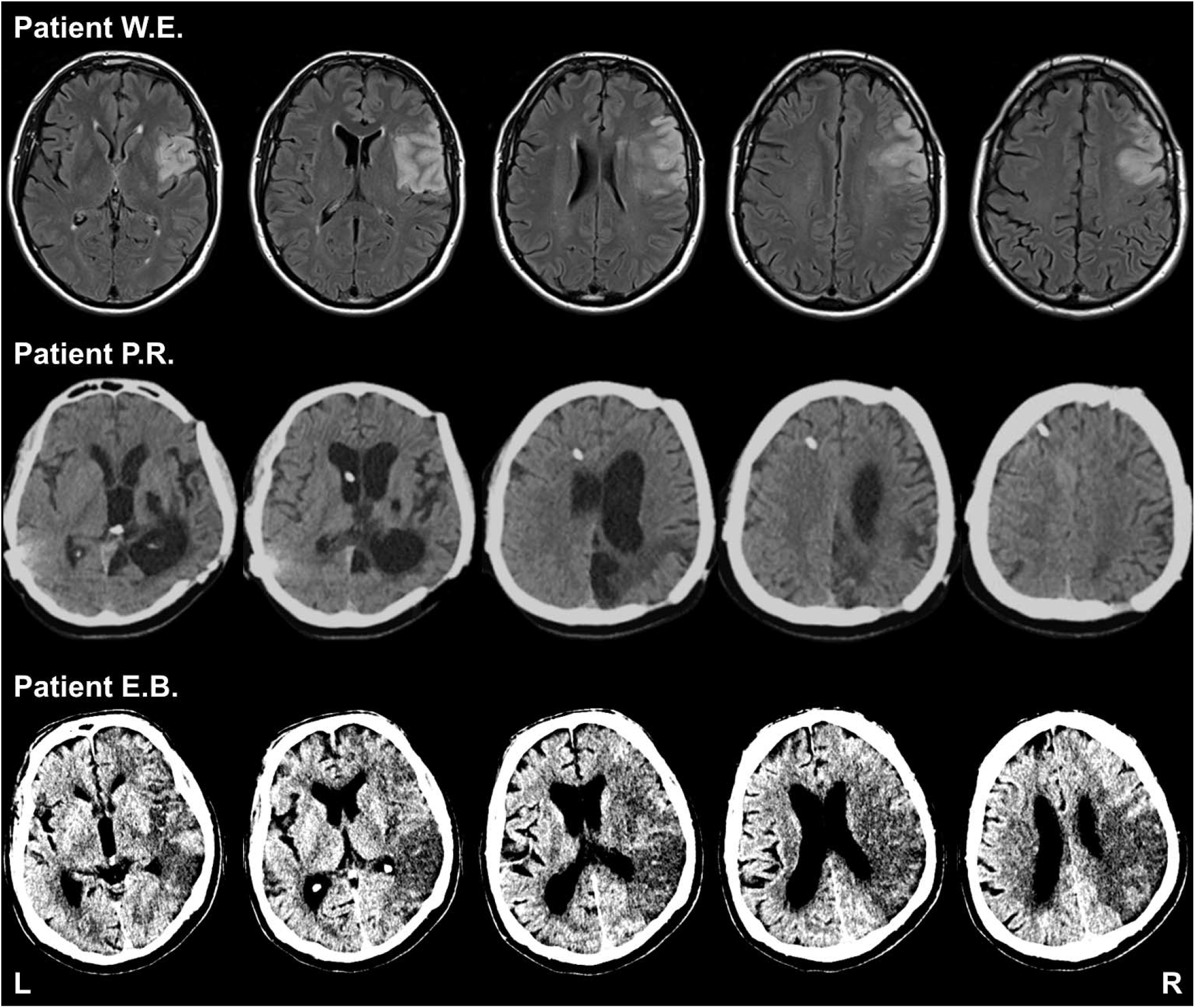
Brain lesions of the patients suffering from spatial neglect.

Patient P.R. is a 35-year-old right handed male patient, who suffered a traumatic brain injury and was found unconscious by his wife 736 days before participating in the present study. Initial CT-imaging at the time of injury uncovered a parieto-temporal epidural hematoma in the right hemisphere. Treatment in the acute phase of the injury included a craniotomy to drain the hematoma, followed by a right-sided decompressive craniectomy. P.R. suffered from tetraparesis, accentuated on the left side, dysarthria, and spatial neglect to the left. A CT scan performed 33 days before participation in the present experiment showed a right-sided lesion affecting medial and lateral parts of parieto-temporal cortex and extending into the right insula and basal ganglia (Fig. 1).

Patient E.B. is a 68-year-old right-handed male patient, who suffered a stroke 104 days prior to his participation in the present study. E.B. initially presented with severe left-sided hemiparesis, dysarthria, and spatial neglect. Initial imaging uncovered an occlusion of the M1 segment of the middle cerebral artery in the right hemisphere, which was treated by mechanical thrombectomy. A CT conducted on the day of symptoms onset uncovered a temporo-parietal infarct in the territory of the right middle cerebral artery (Fig. 1).

The control group consisted of a sample of six healthy subjects (1 male and 5 females; average age 26.3±3.4 years SD). All subjects reported to be right-handed, had normal or corrected to normal vision, and reported no vestibular or neurological deficit. These subjects have been examined in a previous study (Linder et al., 2021). All subjects provided their informed consent according to our institutional ethics board guidelines prior to our experiment.

### Neuropsychological examination

Neuropsychological examination included the following four tests of spatial neglect: the Letter Cancellation Test (Weintraub & Mesulam, 1987), the Bells Test (Gauthier et al., 1989), a Copying Task (Johannsen & Karnath, 2004), and a Line Bisection Task (following the procedure described by McIntosh et al. [2005]). All four neglect tests were performed on a Samsung S7+ tablet with screen dimensions 285x185mm. The severity of spatial neglect in the cancellation tasks was determined by calculating the center of gravity of the target stimuli marked in the search fields, i.e. the Center of Cancellation (CoC; Rorden & Karnath, 2010). A CoC value ≥ 0.08 indicated left-sided spatial neglect (Rorden & Karnath, 2010). The Copying Task consisted of a complex scene consisting of four objects (fence, car, house, tree), points were assigned based on missing details or whole objects. One point was given for a missing detail, two for a whole object. The maximum number of points is therefore 8. A score higher than 1 (i.e. > 12.5% omissions) indicated neglect (Johannsen & Karnath, 2004). In the Line Bisection Task, patients were presented four different line lengths eight times each, i.e., 32 lines in total (McIntosh et al., 2005). The cut-off value for spatial neglect was an endpoint weightings bias (EWB) value ≥ 0.07 (McIntosh et al., 2017).

### Experimental procedure

We used a 3T Siemens MAGNETOM Prisma MRI Scanner to apply MVS. No radio frequency (RF) or gradient coil fields were applied. Contrary to the standard subject positioning in neurology/neuroscience, neglect patients and controls entered the scanner with the feet first (Fig. 2). Therefore, the magnetic field vector of our MRI system pointed from subject’s head to the toes. To this end a 20-channel head coil (Siemens Head/Neck 20 A 3T Tim Coil) was placed at the feet-end of the scanner table. We tried to maximize the effects of MVS on the horizontal canals by positioning subjects with their heads tilted -30° backwards inside the head-coil (canthus-tragus-line vs. vertical; cf. Roberts et al., 2011; Boegle et al., 2016). Cushions were used to additionally stabilize head position inside the coil to prevent head movements. A mirror was mounted on top of the head-coil to provide subjects with an indirect view on our custom-made black visual search-screen. The screen was placed behind the coil on the scanner table at about 110 cm viewing distance. Various glass-fiber-cables penetrated the screen at discrete locations and allowed the presentation of small visual stimuli by feeding these cables with LED light signals. Maximal target distance from the screen center amounted to ±12° visual angle in the horizontal direction and to -5° to 6° in the vertical direction. An IR-video-camera unit was mounted next to the mirror to record a video-signal of subjects’ right eye. The study was conducted in complete darkness by extinguishing all light sources and covering subjects with a black blanket (cf. Fig. 2a). This was important, as any visible stimulus that would be present in the subjects’ visual field could help them to suppress an MVS-induced VOR as well as to explore the visual scene and to perform the SSA task in an unbiased fashion.

**Fig. 2.**
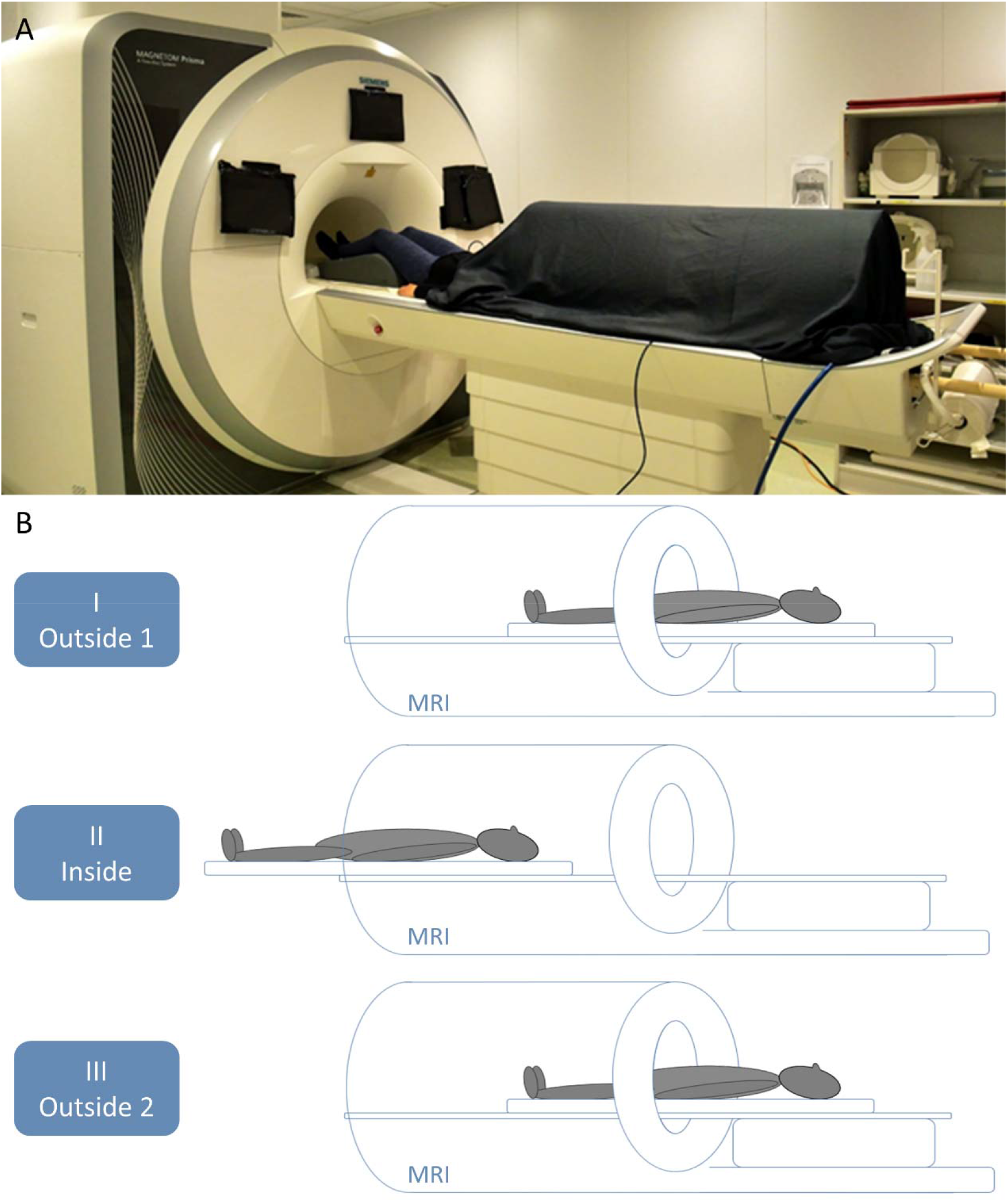
Experimental setting and procedure. **A** Contrary to the standard subject positioning in neurology/neuroscience, neglect patients and controls entered the scanner with the feet first. Subjects were covered with a black blanket extinguishing all light sources. **B** Three consecutive conditions were performed: I. measuring outside the scanner with the patients’ head at its maximal horizontal displacement from the scanner center (“*outside 1*”), II. measuring inside the scanner where the strength of the uniform magnetic field is 3 Tesla (“*inside*”), followed by III. measuring outside the scanner again (“*outside 2*”).

The procedure of the experiment is illustrated in Figure 2B. Three consecutive conditions were performed: measuring outside the scanner (*outside 1* phase) with the patients’ head at its maximal horizontal displacement from the scanner center (head coil at ∼ 215cm/0cm horizontal/vertical displacement); the strength of the magnetic field in this position is roughly 50 times smaller than at the scanner center (head coil at ∼ 10 cm/0cm horizontal/vertical displacement). Subsequently, we measured inside the scanner (*inside* phase) where the strength of the uniform magnetic field is 3 Tesla, followed by again measuring outside the scanner (*outside 2* phase). Each of the three phases started with an initial calibration for eye-tracking. Next, we performed a nystagmus measurement. For this measurement, subjects were instructed to fixate at a dim light-point presented in the center of their visual field for 5 seconds. Then the dim light-point was switched off and subjects were asked to try maintaining central fixation for 60 seconds in otherwise complete darkness. This task was used to quantify subjects’ horizontal VOR during an instructed fixation. Finally, subjects performed a “*visual search task*”. During this task, subjects were asked to find and fixate transient light stimuli. We presented 6 search targets over a period of 5 seconds. Locations of search targets were pseudorandomized across the three phases. The presentation of these stimuli only served to maintain the subjects’ motivation to search for possible targets throughout the search period. Thus, for most of the time (i.e., 140 seconds; total duration of visual search task: 172 seconds), no visible target was present and the subjects were searching in complete darkness. For data analysis we discarded the time periods with targets present (plus an additional grace period of 5 seconds after a light stimulus was turned off; this was done to prevent carry-over effects from prior target-fixation). The search task always ended with a central light stimulus presented during the last 2 seconds (also not considered for data analysis). Subjects’ heads were in the center of the MRI-scanner for approximately 10 minutes. For the *outside 2* phase, we ensured that our experimental measures were only obtained after subjects had left the MRI-scanner for approximately 3 minutes in order to account for putative MVS-aftereffects (Jareonsettasin et al., 2016).

### Stimulus Generation

A WIN10 laptop PC was used to control our experiment through custom scripts using MATLAB R2015b 32bit (MathWorks) in combination with Cogent 2000 and Cogent Graphics (by FIL, ICN, LON at the Wellcome Department of Imaging Neuroscience, University College London) as well as with the MATLAB Support Package for Arduino. Light stimuli were generated through 8 red LEDs (Type: L-513HD; Dropping resistor: 58kΩ) using an Arduino UNO R3-compatible microcontroller (Funduino UNO R3) that was controlled by MATLAB. The intensity of six of those LEDs could be adjusted using the Arduino’s pulse code modulated (PCM) analog voltage output (0V-5V). The remainder of the LEDs could only be switched on/off (0V or 5V). This way we were able to deliver dim light stimuli through the fiber-ends inside the scanner. During the visual search task (see below) the voltage of the 6 search target LEDs was PCM-modulated across the 5 seconds presentation time for a given target, starting with 0.1V while doubling voltage in 1 second steps up to 1.6V. Through this manipulation we slowly increased the visibility of the search targets to increase the likelihood that subjects would find them. For patient E.B., the search targets were always shown with full intensity in order to improve recognizability for him. Apart from these transient light stimuli, subjects remained in complete darkness. The six search targets were at the following x/y–locations (values denote visual angle; positive values represent rightward/upward directions, respectively): -6°/5°; 6°/5°; -6°/-5°; 6°/-5°; -12°/-1°; 12°/1°. The final central search target was always at 0°/0°. Finally, the following 5 targets served for eye-calibration: 0°,0°; -6°/5°; 6°/5°; -6°/-5°; 6°/-5°.

### Eye Tracking

The position of subjects’ right eye was monitored at 50 Hz sampling rate with an MR-compatible camera with integrated infrared LED illumination (MRC Systems; Model: 12M-i IR-LED). We used the ViewPoint Monocular Integrator System and the View Point Software (Arrington Research, software version 2.8.3.437) to digitize the eye-camera video and to obtain uncalibrated eye-position data (by means of dark pupil tracking). Eye-tracking was realized on a different WIN PC that was remote-controlled through the ViewPoint Ethernet-Client running on our laptop PC. Eye movement analyses were performed off-line using custom routines written in MATLAB R2017b (MathWorks). Details are given in Lindner et al. (2021).

### Statistical Analyses

The statistical procedures to test for any MVS-effects on the behavior of our group of healthy control subjects have been described in detail before (Lindner et al., 2021). In order to probe for comparable MVS-induced effects on the horizontal VOR as well as the horizontal center of visual search in our group of neglect patients, we took individual samples of de-saccaded horizontal eye velocity and horizontal saccade endpoints of the respective experimental phases as measures. Since these latter data were not normally distributed in all cases (Shapiro-Wilk-Test; p<0.01), we performed Wilcoxon rank sum tests throughout (two-tailed tests: *outside 1* vs. *inside* and *outside 2* vs. *inside*). To compare the neglect patients’ initial search behavior to that of the control group, we conducted a two-tailed Wilcoxon rank sum test for *outside 1*.

## Results

Figure 3 illustrates the eye-data of a single patient with spatial neglect throughout all tasks and phases. During the *inside* phase we observed the typical saw-tooth-like eye movement pattern, which is characteristic for a vestibular nystagmus (Fig. 3a, left column). It consisted of a slow leftward VOR that was accompanied by fast resetting saccades in the opposite rightward direction. This prototypical nystagmus-pattern was largely reduced for the *outside 1&2* phases. Figure 3b demonstrates a 2D-plot of the patient’s eye position during search, including only those time epochs when no target stimulus was presented (cf. methods section). As was expected for patients with spatial neglect (Karnath et al., 1996, 1998), we observed a biased ocular exploratory search pattern towards the ipsilesional right side during the *outside 1* phase (cf. Fig. 3b). This pathological search bias was clearly reduced when the patient was *inside* the scanner, and increased again during the *outside 2* phase.

**Fig. 3.**
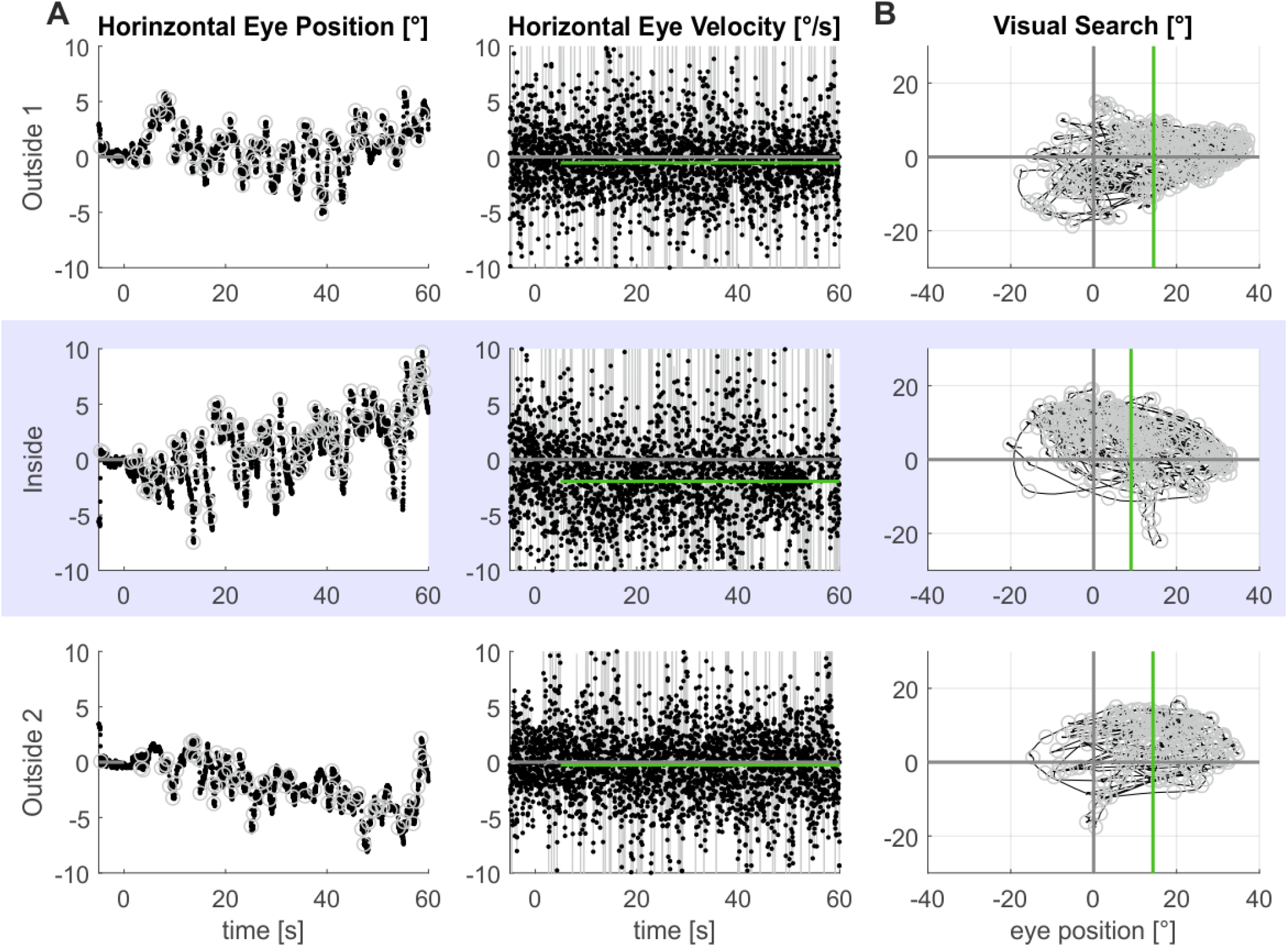
MVS-induced oculomotor behavior in one exemplary patient with spatial neglect (patient W.E.) in the three phases (*outside 1, inside, outside 2*). **A** shows data from the nystagmus measurement. Horizontal eye position data are shown in the left column. Saccade endpoints are depicted by the grey circles. Corresponding horizontal eye velocity traces are shown in the middle column. Note that the grey peaks in these time-courses refer to individual saccades, which have been removed from the eye velocity records to allow estimation of slow-phase velocity in isolation (the green lines indicate the respective horizontal VOR-estimates). **B** illustrates the 2D eye-position data from the visual search task (time periods with search targets plus 5 seconds were excluded). The horizontal center of visual search is depicted by the green vertical lines, reflecting the mean of horizontal saccade endpoints (grey circles). Positive values indicate the rightward/upward direction.

The same principle pattern of results was obtained in all of our neglect patients. Horizontal eye velocity increased significantly towards the leftward direction in the *inside* condition as compared to both *outside* conditions (all p<0.001; two-tailed Wilcoxon rank sum tests; see Fig. 4). This demonstrates that an MVS-induced VOR was present in all three cases.

**Fig. 4.**
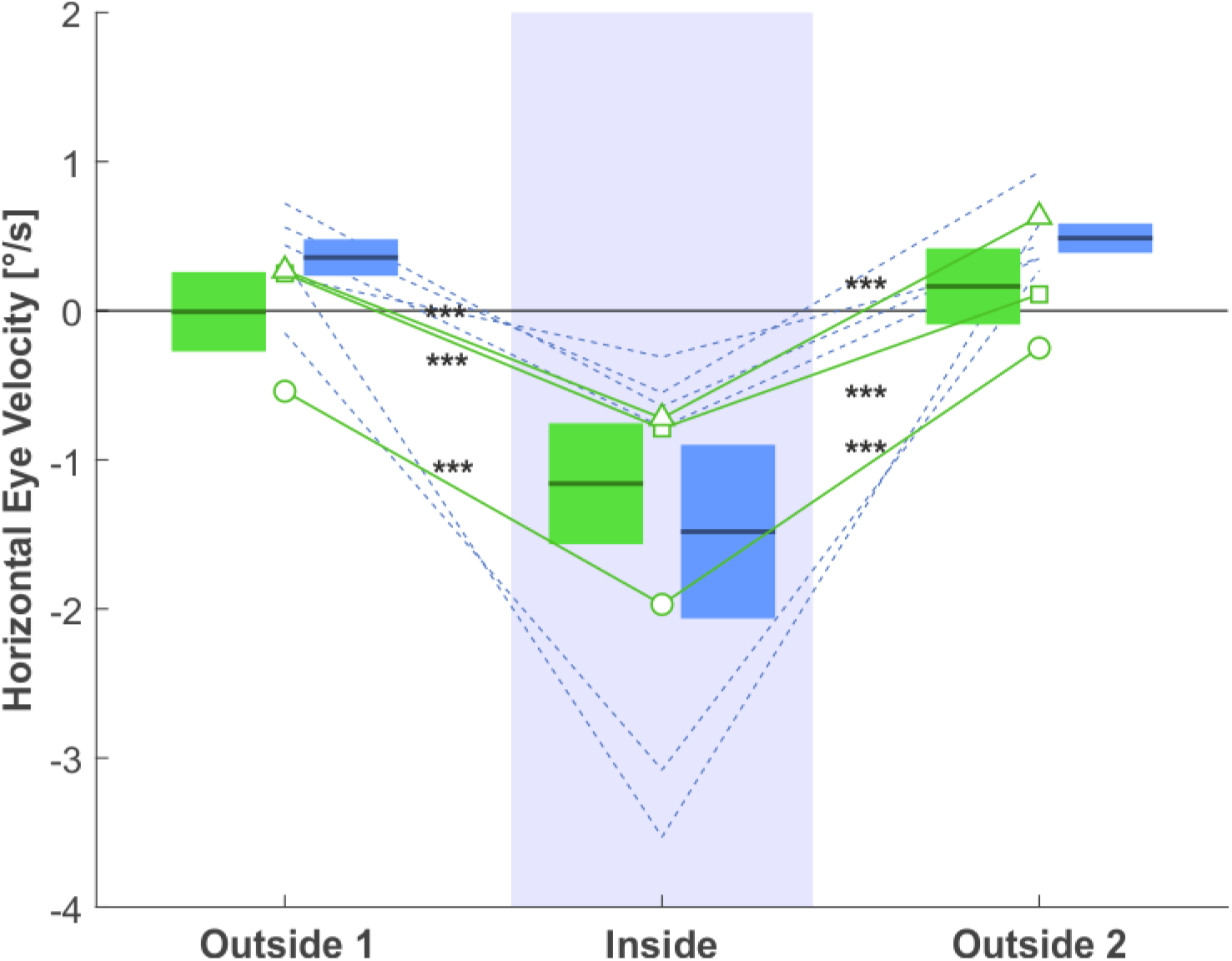
Horizontal eye velocity. Neglect patients’ means and standard errors are depicted as green boxes and green lines, respectively. As a reference, we additionally show respective data for six healthy subjects from our earlier study in blue (cf. Lindner et al., 2021). Also, the medians of the de-saccaded horizontal eye-velocity traces is shown for individual patients (green symbols: circle, square, and triangle refer to the individual neglect patients). There was a significant increase in leftward eye velocity for the *inside* condition in all cases (*** p<0.001; two-tailed Wilcoxon rank sum tests).

In terms of their visual search performance, patients initially clearly deviated from the healthy subjects’ measures in our earlier study (Lindner et al., 2021): As to be expected for spatial neglect, patients’ horizontal mean of visual search was significantly shifted towards the right during *outside 1* (p<0.028; two-tailed Wilcoxon rank sum test; cf. Fig. 5). More importantly, patients’ horizontal mean of visual search shifted significantly towards the left during the *inside* phase and thus reduced the neglected part of space. This was true for each individual neglect patient (all p<0.001; two-tailed Wilcoxon rank sum tests). The reduction of spatial neglect (as quantified through visual search) was only temporary in patient W.E. as search performance shifted back towards the right from *inside* to *outside 2* (p<0.001; two-tailed Wilcoxon rank sum test). Interestingly, patient P.R. maintained the reduction of the rightward bias in search performance also during *outside 2* (p=0.61), while patient E.B.’s search performance did even exhibit further improvement from *inside* to *outside 2* (p<0.001).

**Fig. 5.**
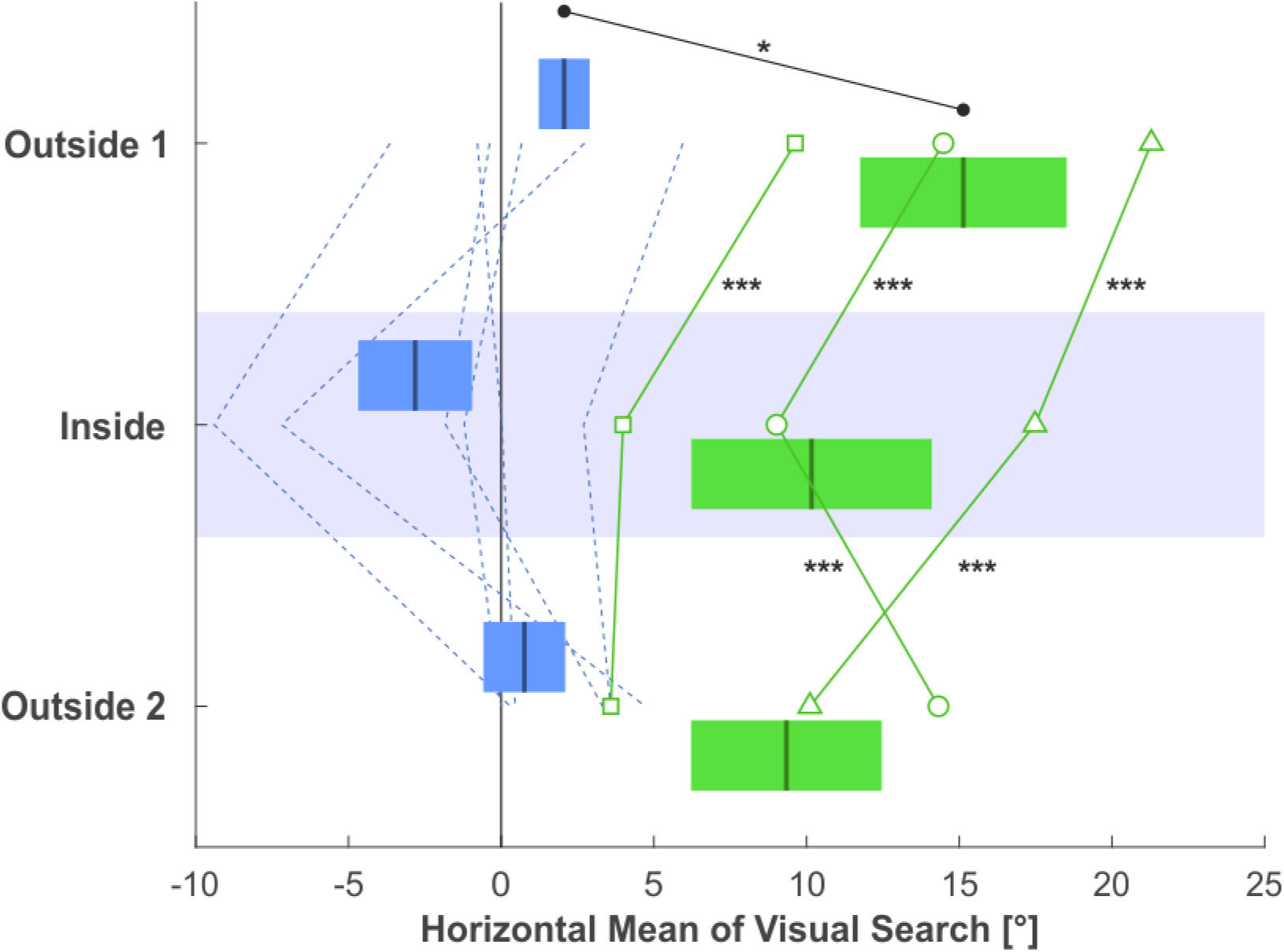
Horizontal means of visual search behavior. Neglect patients’ means and standard errors are depicted as green boxes and green lines, respectively. As a reference, we additionally show respective data for six healthy subjects from our earlier study in blue (c.f. Lindner et al., 2021). Note that, as to be expected for spatial neglect, patients mean of visual search was initially shifted to the right as compared to healthy subjects during *outside 1* (* p<0.028; two-tailed Wilcoxon rank sum test). Also, the horizontal means of visual search is depicted for individual patients (green symbols: circle, square, and triangle refer to the individual neglect patients). There was a significant leftward shift from *outside 1* to *inside* in all cases (*** p<0.001; two-tailed Wilcoxon rank sum tests).

## Discussion

The present findings demonstrate that the bias of spatial attention and exploration in patients with spatial neglect is indeed reduced by MVS. Lying in a 3T scanner ameliorated the patients’ ipsilesionally biased distribution of overt attention and the corresponding neglect on the left of the search-space.

The present results closely resemble the known effects of caloric vestibular stimulation (CVS) on spatial exploration and orientation in neglect patients. When their left external auditory canal was irrigated with cold water, stroke patients with spatial neglect showed a reduction of their neglect of contralesionally located stimuli (Rubens, 1985; for review cf. Rossetti & Rode, 2002) as well as of their biased distribution of overt attention in visual search (Karnath et al., 1996). Moreover, it has been demonstrated that the typical bias in subjective straight-ahead body orientation in neglect patients is reduced by CVS (Karnath, 1994). Conversely, in healthy subjects CVS in darkness induces a bias in subjective straight-ahead (SSA), mimicking the bias of the SSA in neglect patients towards their ipsilesional side (e.g. Karnath, 1994; Chokron & Imbert, 1995; Kapoor et al., 2001). Finally, CVS biases healthy subjects’ scan path during visual search (Karnath et al., 1996). When exploring their surroundings for possible targets, subjects’ eye movements are no longer symmetrically distributed in the horizontal dimension but biased towards the side of cold CVS. To summarize, the present results in a consecutively admitted group of patient with spatial neglect as well as the previous observations in healthy subjects by Lindner et al. (2021) demonstrate a close physiological correspondence of MVS and CVS. The two types of vestibular stimulation evoke very similar if not identical behavioral effects in stroke patients with spatial neglect.

However, regarding the use of vestibular stimulation in the rehabilitation of spatial neglect, MVS is far superior to CVS. While CVS is highly unpleasant due to the application of cold water to the ear, the physiological effect of MVS at 3T does not evoke any unpleasant sensations; actually, it is not even directly perceived while the subject lies in the scanner (Milian et al., 2013). More importantly, the effect of CVS (in healthy subjects as well as in patients with neglect) lasts only a few minutes, while the physiological effect of MVS persists for the entire time while the subject lies in the scanner (Roberts et al., 2011; Jareonsettasin et al., 2016; for review see Ertl & Boegle, 2019). So far, this has been shown for a time period of 90 minutes inside a 7T scanner. During this time, the horizontal nystagmus diminished but persisted and did not extinguish (Jareonsettasin et al., 2016). Thus, for therapeutic considerations in spatial neglect, MVS is the much more appropriate type of vestibular stimulation. The tonic nature of MVS could serve as a noninvasive and comfortable means to continuously stimulate the labyrinth of stroke patients with spatial neglect. However, whether MVS is actually suitable for reducing neglect symptoms in a therapeutic sense, i.e., whether it might help to induce longer-term or even lasting plastic changes in the patients’ pathologically altered spatial representations, still remains to be proven in future studies. Yet, the fact that the MVS-induced reduction of the exploration bias was maintained in one neglect patient and even further reduced in another neglect patient also after they had left the scanner seems promising. Finally, the availability and exclusion criteria of MRI as well as the patient’s potential burden associated with lying in the narrow tube of an MRI scanner must also be weighed against any potential therapeutic benefit. Nevertheless, the present study supports the idea that an intervention as simple as lying in a 3T MRI scanner helps reduce spatial neglect through MVS.

## Acknowledgements

This work was supported by the Deutsche Forschungsgemeinschaft.

## Notes

### Competing Interest Statement

The authors have declared no competing interest.

### Summary of Updates

Illustration of procedures in Figure 2B was optimized.

